# Allogeneic gene-edited HIV-specific CAR-T cells secreting PD-1 blocking scFv enhance anti-HIV activity in vitro

**DOI:** 10.1101/2022.01.14.476413

**Authors:** Hanyu Pan, Jing Wang, Huitong Liang, Zhengtao Jiang, Lin Zhao, Yanan Wang, Xinyi Yang, Zhiming Liang, Xiaoting Shen, Qinru Lin, Yue Liang, Jinglong Yang, Panpan Lu, Yuqi Zhu, Weirong Xiang, Min Li, Pengfei Wang, Huanzhang Zhu

## Abstract

HIV-specific chimeric antigen receptor (CAR) T cells have been developed to target latently infected CD4+ T cells that express virus either spontaneously or after intentional latency reversal. However, the T-cell exhaustion and the patient-specific autologous paradigm of CAR-T hurdled the clinical application. Here, we created HIV-specific CAR-T cells using human peripheral blood mononuclear cells and a 3BNC117-E27 CAR (3BE CAR) construct that enables the expression of PD-1 blocking scFv E27 and the single-chain variable fragment of the HIV-1-specific broadly neutralizing antibody 3BNC117 to target native HIV envelope glycoprotein (Env). In comparison with T cells expressing 3BNC117-CAR alone, 3BE CAR-T cells showed greater anti-HIV potency with stronger proliferation capability, higher killing efficiency (up to ~75%) and enhanced cytokine secretion in the presence of HIV envelope glycoprotein-expressing cells. Furthermore, our approach achieved high levels (over 97%) of the TCR-deficient 3BE CAR-T cells with the functional inactivation of endogenous TCR to avoid graft-versus-host disease without compromising their antiviral activity relative to standard anti-HIV CAR-T cells. These data suggest that we have provided a feasible approach to large-scale generation of “off-the-shelf” anti-HIV CAR-T cells in combination with antibody therapy of PD-1 blockade, which can be a powerful therapeutic candidate for the functional cure of HIV.

## Introduction

Since acquired immunodeficiency syndrome (AIDS) was first reported in 1981,^1^ 36.3 million people have died from AIDS-related illness.^2^ Although the highly effective combined antiretroviral therapy (cART) can suppress virus loads to undetectable levels in infected individuals, it does not eliminate the viral reservoirs and needs lifelong adherence to expensive regimens with potential toxic effects.^3^ New approaches to achieve a HIV cure and avoid the burden of lifelong ART are urgently needed.^4^ For the past few years, one of the broadly defined categories of a cure for HIV infection, a functional cure, has become a global research priority.^3,5^ As is essential to boost the immune response in infected individuals so that antiviral drugs can be discontinued,^6^ a successful strategy for a functional cure will require potent and persistent cellular immune surveillance and the adoptive transfer of effector T cells modified with a chimeric antigen receptor (CAR) might be an applicable approach.^7^ CAR-T therapy, which provide non-MHC–restricted recognition of cell surface components to kill the target cells with high efficiency,^8^ has achieved tremendous successes in the treatment of hematological malignancies.^9^ Notably, one of the earliest clinical trials of CAR-T therapy was for the treatment of AIDS in 1994, using the “first generation” CD4-based CAR constructs.^10^ Unfortunately, those CAR-T cells showed no clear benefits in HIV-infected individuals, despite of the safety^11^ and decade-long persistence *in vivo*.^12^ In the wake of the appearance of the “second generation” CAR containing costimulation modulates such as CD28 and 4-1BB,^13^ the efficacy of anti-tumor CAR-T cells was significantly improved.^14,15^ The success obtained with CD19 CAR-T cells rekindled interest in anti-HIV CAR-T.

Since the CD4 domain might allowed HIV-1 infection of CAR-T cells and there could be selection for viral escape through reduced CD4 binding,^16^ several groups have tried to manufacture new anti-HIV CAR-T by replacing the extracellular CD4+ domain of the CAR with abroadly neutralizing antibody (bNAb) single-chain variable fragments (scFv), such as VRC01, 3BNC117 and 10E8.^17,18,19,20^ The safety, tolerability, and therapeutic efficacy of 3BNC117 in HIV-infected human individuals have been confirmed by Phase I and Phase IIa clinical trials.^21,22^ Previous studies have also shown the potent antiviral activity of 3BNC117-based CAR *in vitro*^17^ and *in vivo*^23^. However, the immunosuppression might be a huge obstacle to the clinical application of bNAb-based CARs. The inhibitory receptor programmed death 1 (PD-1) has been reported to be markedly upregulated on the surface of HIV-specific CD8 T cells *in vivo*, which resulted in the reduced capacity of those cytotoxic T lymphocytes (CTLs) to produce cytokines as well as an impaired capacity to proliferate.^24,25,26^ On the other hand, there has been increasing evidence that PD ligand 1 (PD-L1) expression was induced on a variety of cell types by chronic HIV infection. ^27,28,29,30^ Previous studies have demonstrated that the blockade of PD-1 by using monoclonal antibodies enhanced T cell cycling and differentiation and resulted in rapid expansion of virus-specific CD8 T cells with improved functional quality in simian immunodeficiency virus (SIV)-infected rhesus macaques.^31,32^ Those improved immune responses by PD-1 blockade induced significant reductions in plasma viral load and prolonged the survival of SIV-infected macaques.^31^ The PD-1 blockade could be further combined with the blockade of cytotoxic T lymphocyte-associated protein 4 (CTLA-4), which could induce robust viral reactivation in plasma and peripheral blood mononuclear cells,^32^ which could make sense for the “shock and kill” strategy.

Another challenge to developing anti-HIV CAR-T therapy for clinical application comes from the patient-specific autologous paradigm of CAR-T. The majority of current CAR-T clinical trials utilize autologous T cells from each patient, which entail unwarranted delays inherent to the generation of therapeutic products with prohibitive cost. With respect to HIV-infected individuals, it can be much more difficult to obtain enough T cells of good quality for manufacturing therapeutic products since their immune system has been seriously damaged. Recent years, many researchers have been paying attention to “off-the-shelf” CAR-T. Torikai et al. knockout T-cell receptor (TCR) αβ expression from allogeneic CD19-CAR T cells by zinc finger nucleases (ZFNs).^33^ Similarly, Poirot et al. manufactured the third-party TCR-deficient CD19-CAR T cells via the application of transcription activator–like effector nuclease (TALEN) gene-editing technology and avoided graft-versus-host disease (GvHD) in the xenogeneic mouse model.^34^ Furthermore, Eyquem et al. directed a CD19-CAR to the TCR α constant (*TRAC*) locus by using homology-directed recombination (HDR) together with CRISPR/Cas9 technology, and demonstrated great anti-tumor potency of the TCR-negative CAR-T cells.^35^

Here, we designed and produced anti-HIV CAR-T cells derived from 3BNC117 targeting HIV-1 envelope glycoprotein as well as secreting PD-1 blocking scFv E27,^36^ and then disrupted the expression of endogenous TCR in those CAR-T cells to avoid GvHD. We assessed abilities of the 3BNC117-E27/TCR-deficient CAR-T cells to eliminate HIV target cells, and our results showed that those edited cells acquired enhanced T-cell potency vastly outperformed conventional 3BNC117-CAR T cells and could be a potential candidate for the functional cure of HIV.

## Results

### The genetic construction and expression of 3BNC117-based CARs against HIV-1

The whole composition of 3BNC117 CAR (3B CAR), which consist of the 3BNC117 scFv, IgG4 hinge, CD8 TM and the 4-1BB signaling domain fused to the CD3 ζ signaling domain, was cloned to the pTRPE lentiviral vector to generate a “second generation” CAR vector (figure 1A). As it was reported that PD-1 expression on HIV-specific T cells was associated with T-cell exhaustion and disease progression,^24^ we additionally modified the CAR construct to include a HA-tagged anti-human PD-1 scFv E27 (3BNC117-E27 CAR (3BE CAR), figure 1A), which had been reported to block the PD-1/PD-L1 pathway by competitively binding to PD-1 and keep CAR-T cells from exhausting^36^ (figure 1B).

**Figure 1.**
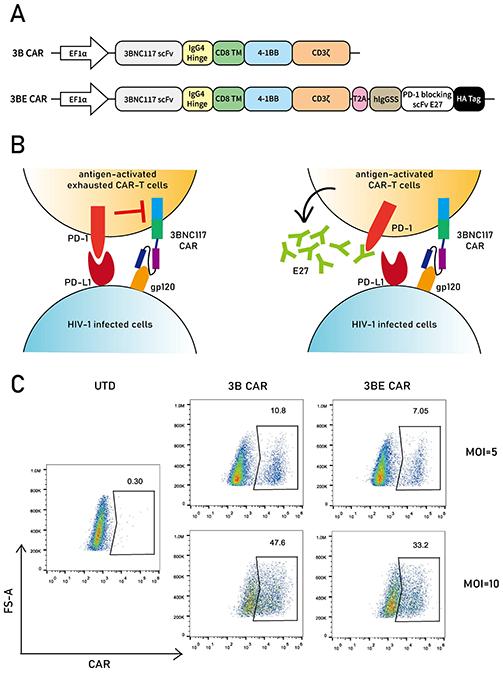
Design and expression of 3BNC117-based CAR constructs. **(A)** Schematic of vectors encoding the 3BNC117 CAR (3B CAR) and 3BNC117-E27 CAR (3BE CAR). EF1α indicated in the diagram is promoter sequences. **(B)** Schematic representation of the way that E27 blocking the binding of PD-1 with PD-L1 to keep CAR-T cells from exhausting. **(C)** Representative flow plot showing surface CAR expression on CD3+ T cells by detecting human Fab through flow cytometry. UTD cells served as a negative control.

The CAR genes were delivered by lentiviral vectors to Primary CD3+ T lymphocytes isolated from human peripheral blood mononuclear cells (PBMCs) to manufacture anti-HIV CAR-T cells, and flow cytometry using a goat antibody against human antigen-binding antibody fragment (Fab) was performed on 3 days after transduction to demonstrate the expression of each CAR on the cell surface. At MOI ~5, the transduction of CAR lentiviral vectors produced about 7%-10% positive cells; while at a higher MOI (~10), the frequency of CAR-positive cells could be increased to about 30%-50% (figure 1C).

### Function of PD-1 blocking scFv E27 secreted by CAR-T cells *in vitro*

The expression and secretion of PD-1 blocking scFv E27 by 3BE CAR-T cells were demonstrated by western blot analysis detected with anti-HA-tag antibody (figure 2A). It was showed that E27 existed more in the supernatant than in the cytoplasm, indicated the high efficiency of the secretion signal we chose. To investigate the anti-HIV efficacy of E27-secreting CAR-T cells *in vitro*, we co-cultured 3BE CAR-T cells with LHL2-3, which had PD-L1 overexpression by lentivirus based on HL2/3^37^ and could constitutively express Env and PD-L1 at the cell surface,^23^ at different ratio and tested the specific cytotoxicity using the LDH release assay. The results suggested that both groups of 3BNC117 CAR-T cells efficiently eliminated Env+ cells at 10:1 ratio, while CAR-T cells secreting E27 scFv showed greater potency (figure 2B).

**Figure 2.**
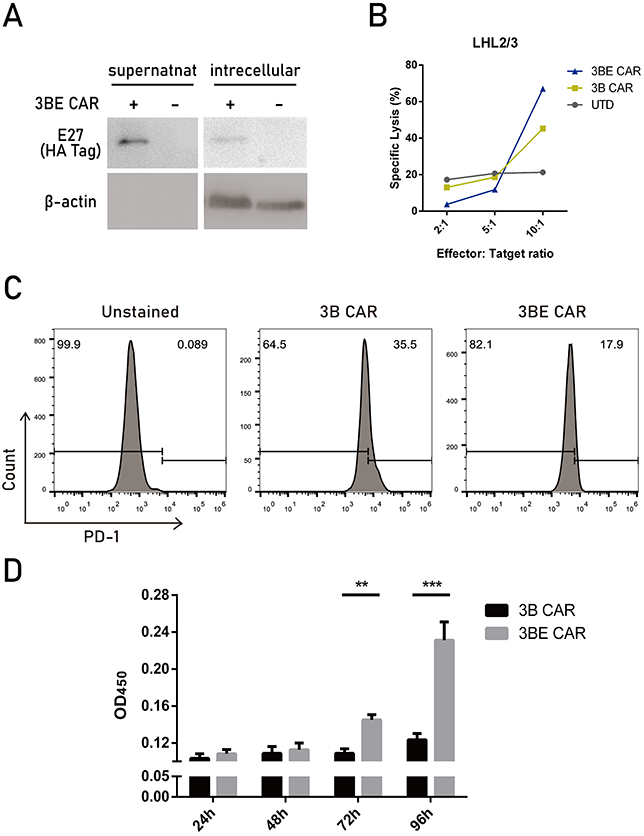
The expression and secretion of PD-1 blocking antibody E27 enhances the proliferation and killing ability of CAR-T cells incubated with target cells. **(A)** Western blot on supernatant and intracellular material from cells transfected to express the secretable scFv with the CAR, detected with anti-HA-tag antibody. **(B)** Anti-HIV CAR-T cells were incubated with LHL2/3 cells at different ratios (2:1; 5:1; and 10:1) for 4 h and direct killing of LHL2/3 was performed using the LDH release assay. **(C)** Quantification of PD-1 detection by flow cytometry on 3BE CAR-T cell, as compared to 3B CAR-T cells. **(D)** Anti-HIV CAR-T cells were incubated with LHL2/3 cells at 1:1 for 24 h to 96 h, and proliferation of CAR-T cells was detected by CCK-8. Data shown is representative of 3 independent experiments. ^**^P < 0.01, ^***^P < 0.001

Since E27 had been reported as a PD-1 blocking agent, we evaluated binding of secreted E27 scFv to PD-1. The results of flow cytometry showed that 3BE CAR-T cells significantly decreased surface detection of PD-1 compared to the cells modified to express the CAR alone (3B CAR-T cells) (figure 2C), which was consistent with previous reports^36^ and suggested that E27 binds in an autocrine manner to secreting CAR-T cells.

As it has been confirmed that PD-1/PD-L1 pathway played an important role in T cell exhaustion, anergy, and/or apoptosis,^38,39,40,41^ we hypothesized that PD-1-blocking scFv E27 could overcome the suppression of anti-HIV CAR-T cells. To verify this, 3B or 3BE CAR-T cells were co-cultured with LHL2/3 at 1:1 ratio for 24 hours to 96 hours, and proliferation of those CAR-T cells was detected by Cell Counting Kit-8 (CCK-8). Consistent with our assumption, 3BE CAR-T cells showed significantly stronger proliferation capability compared to 3B CAR-T cells (figure 2D), providing evidence that E27 provide a proliferative advantage of anti-HIV CAR-T cells in the presence of target cells.

### Disruption of human *TRAC* gene and functional inactivation of TCR in primary T cells with high efficiency by CRISPR/Cas9

To manufacture "Off-the-Shelf" anti-HIV CAR-T cells unable to initiate xenogeneic GvHD, the key is efficient genomic editing to generate gene-disrupted T cells that are deficient in TCR. For this purpose, we first selected four sgRNA-targeting sites on *TRAC* according to previous published results,^35,42,43,44^ all of which are within the first exon of the gene (figure 3A). Cas9 ribonucleoproteins (RNPs), complexes of spCas9 protein with synthetic single guide RNAs (sgRNAs) mentioned above, were electroporated into HEK293T cells and their activity was validated by a mismatch-selective T7E1 surveyor nuclease assay. The results indicated that all the sgRNAs leaded to distinctly detectable cleavage at the genomic loci of *TRAC* except sgRNA-1, and it was confirmed a higher cleavage rate with sgRNA-4 which we used in further experiments (figure 3B).

**Figure 3.**
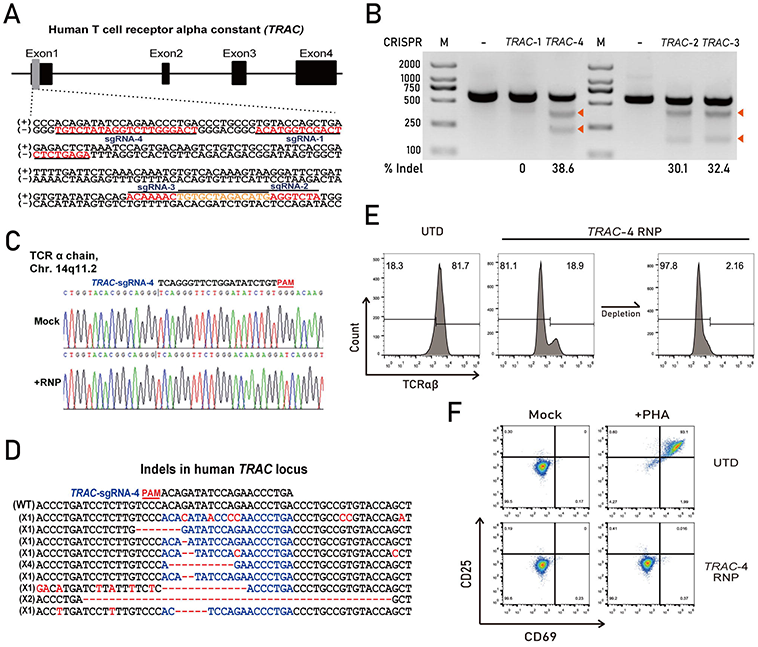
CRISPR/Cas9 mediates efficient TCR disruption in T cells. **(A)** Schematic diagram of sgRNA-targeting sites on *TRAC*. All the targeting sites are within exon1 of the gene. **(B)** RNPs targeting *TRAC* locus were electroporated into HEK293T cells. And the amount of TCR-targeted gene disruption were measured by a mismatch-selective T7E1 surveyor nuclease assay on DNA amplified from the cells. The calculated amount of targeted gene disruption in *TRAC* is shown at the bottom. Arrows indicate expected bands. **(C)** Multiple peaks in the Sanger sequencing results show the CRISPR-mediated events of NHEJ at the *TRAC* genomic locus in CD3+ T cells. **(D)** Indels and insertions observed by clonal sequence analysis of PCR amplicons from CD3+ T cells after CRISPR-mediated recombination of the *TRAC* locus. **(E)** TCRαβ expression on CAR-T cells electroporated with *TRAC*-4 RNP and purified by negative selection using microbeads. **(F)** TCR-deficient cells are resistant to TCR stimulation. UTD cells or T cells electroporated with *TRAC*-4 RNP were treated for 36 h with 1 mg/mL PHA and analyzed by flow cytometry for CD25/CD69 expression.

Next, the electroporation of *TRAC*-4 RNP was performed on human primary CD3+ T cells and disruption of the gene was also confirmed by a T7E1 assay (data not shown). Then we amplified and subcloned the target region from the *TRAC*-4 RNP treated T cells. The genomic reading frame was confirmed to shift downstream of the *TRAC* target site by multiple peaks in the Sanger sequencing data flanking the target site showed in figure 3C. And Sanger sequencing data also confirmed that insertions or deletions (indels) caused by NHEJ repair occurred in majority (13/15, 86.7%) of the clones of *TRAC* PCR products, in which all mutations recovered happened precisely at the sgRNA-targeting region (figure 3D).

Even though we had achieved high levels of TCR-negative knockout (TCR-) T cells (about 80% detected by flow cytometry) by utilizing *TRAC*-4 RNP, it would be still necessary to achieve a higher proportion of T cells deficient in TCR to avoid GvHD as much as possible. To this end, a single step of CD3 negative selection was applied to enriched the TCR disrupted population, and the frequency of TCR-cells could be increased to over 97% (figure 3E). For thorough testify the functional inactivation of the TCR gene and the prevention of responses to TCR stimulation in T cells after the electroporation of *TRAC*-4 RNP, the enriched cells mentioned above were treated with phytohemagglutinin (PHA), and non-electroporated T cells served as a positive control. The results showed that TCR-negative cells failed to upregulate the activation markers CD69 and CD25 upon PHA-mediated TCR stimulation (figure 3F), remarkably distinguished from non-electroporated T cells.

### 3BE CAR-T cells have enhanced anti-HIV potency regardless of the deficiency of TCR

TCR-deficient anti-HIV CAR-T cells secreting PD-1 blocking scFv E27 were manufactured by combing the lentiviral transduction of 3BE CAR with the electroporation of *TRAC*-4 RNP.

To compare the anti-HIV potency of those CAR-T cells, we co-cultured LHL2/3 with 3B, 3BE or 3BE/TCR-CAR-T cells, or untransduced CD3+ T (UTD) cells as a control, at different ratio for 8 hours and then assessed the specific killing of those Env+ cells. All the three groups of anti-HIV CAR-T cells displayed robust cytotoxicity against LHL2/3 compared with UTD cells, especially at 10:1 E:T (effector: target) ratio. Dose-dependent effects of those CAR-T cells on the specific cytotoxic activity against Env+ cells were observed in LHL2/3 at the indicated E:T ratios. As shown in Figure 4A, the specific killing of LHL2/3 rose to 1.5-2 folds as the E:T ratio increased from 1:1 to 5:1, and continued to rise to over 2.6 folds at 10:1 E:T ratio. It was worth noting that 3BE CAR-T cells showed stronger killing activity than 3B CAR-T cells at each individual E: T ratio. In addition, it was indicated that TCR disruption had no discernable effect on the cytotoxicity of anti-HIV CAR T cells against LHL2/3 (figure 4A). Furthermore, the expression level of luciferase, which was delivered into LHL2/3 as a reporter gene, was also detected after the coincubation at different E:T ratio to confirm the killing efficiency. The results were quite consistent with what we mentioned above. At 10:1 E:T ratio, the reduction of luciferase expression caused by 3BE CAR-T cells was more remarkable than that caused by 3B CAR-T cells, but was equivalent to that caused by 3BE/TCR-CAR-T cells (figure 4B).

**Figure 4.**
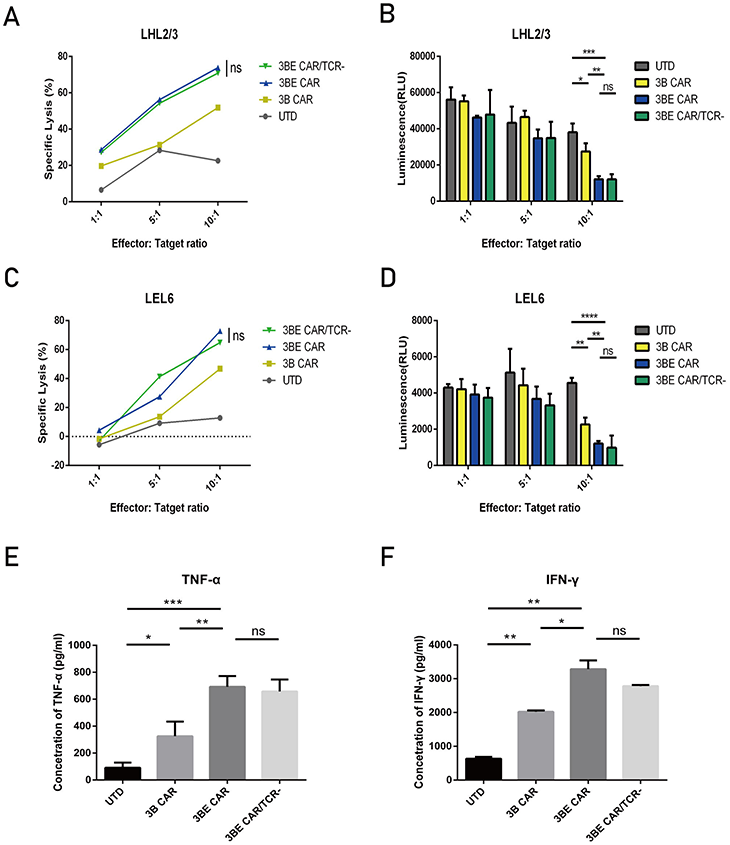
3BE CAR-T cells shows enhanced cytolytic and cytokine secreted function on target cells regardless of the deficiency of TCR. Anti-HIV CAR-T cells were incubated with target cells at different ratios (1:1; 5:1; and 10:1) for 8 h. Direct killing of LHL2/3 cells **(A)**or LEL6 cells **(B)**was performed using the LDH release assay. Detection of luminescence (RLU) in co-cultures reflect the lysis of LHL2/3 cells **(C)**or LEL6 cells **(D)**. Anti-HIV CAR-T cells were co-cultured with LEL6 cells (1 × 10^4^ cells) at 10:1 for 24 h, and supernatants were collected for ELISA to show the production of TNF-α **(E)**and IFN-γ **(F)**in co-cultures. ns p > 0.05, *P < 0.05, **P < 0.01, ***P < 0.001.

Since LHL2/3 derived from Hela cells rather than human T cells, the LEL6 cell line, a luciferase-expressing Env+/PD-L1+ clonal derived from Jurkat cells which had been previously constructed in our laboratory,^23^ was further used in order to examine whether similar results could be obtained in other target cells. The results from these cells also indicated the greater cytotoxic activity of 3BE CAR-T cells than 3B CAR-T cells, even with the deficiency of TCR (figure 4C, D).

To further analyze the effect of the secretion of E27 and the disruption of *TRAC* on CAR-T cell function, the cytokine secretion ability of each kind of anti-HIV CAR-T cells was tested when cultured in the presence of antigen. After co-culture with LEL6 at 10:1 E:T ratio, all the three groups of anti-HIV CAR-T cells produced and released much more TNF-α than UTD cells (Figure 4E), and 3BE CAR-T cells showed enhanced cytokine secretion compared to 3B CAR-T cells. Importantly, 3BE/TCR-CAR-T cells did not demonstrate significantly reduced TNF-α release compared with 3BE CAR-T cells (Figure 4E), indicating that the lack of endogenous TCR expression does not impair the effector function. Similar results were found for the secretion of IFN-γ as well (Figure 4F).

### The secretion of E27 and the deficiency of TCR do not impact the phenotype of CAR-T cells

Since it has been reported that the phenotype of CAR-T cells could impact their function and proliferation *in vivo*,^45,46,47,48,49^ we measured the phenotype of our anti-HIV CAR-T cells 12 days after the CD3/CD28 stimulation.

In comparison with UTD cells, there was no significant difference among the CD62L expression of 3B, 3BE and 3BE/TCR-CAR-T cells (figure 5A). The data showed that the CD62L+ cells in all the groups of anti-HIV CAR-T cells we manufactured were over 60%, which indicated that our CAR-T cells were mainly composed of naive and central memory cells, a phenotype associated with greater *in vivo* anti-HIV activity^35,50^ (figure 5A). The expression of CD45RA was also tested and the percentages of CD45RA+ cells were lower than 7% in all the anti-HIV CAR-T cells (figure 5A). The low level of CD45RA expression reflected that the proportion of terminal effector cells was not large, which suggested a potential advantage in persistence and efficacy *in vivo*.

**Figure 5.**
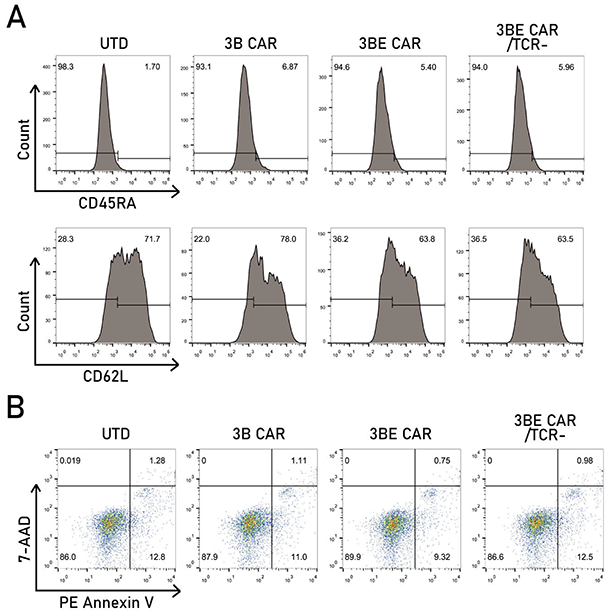
The secretion of E27 and the deficiency of TCR do not impact the phenotype of CAR-T cells. **(A)** The expression of CD45RA and CD62L on UTD,3B CAR 3BE CAR and 3BE CAR/TCR-T cells. **(B)** Apoptosis of UTD,3B CAR 3BE CAR and 3BE CAR/TCR-T cells detected by Annexin V-PE apoptosis detection kit.

In order to measure the toxicity of the delivery of CARs and *TRAC*-4 RNP, the apoptosis of the anti-HIV CAR-T cells was detected by Annexin V-PE apoptosis detection kit. As shown in figure 5B, our anti-HIV CAR-T cells were indistinguishable from UTD cells in terms of the degree of T-cell apoptosis.

According to the results above, we concluded that the secretion of E27 and the deficiency of TCR did not significantly alter the phenotype of anti-HIV CAR-T cells.

## Discussion

Due to the persistence of latent reservoir, which has been a major obstacle to HIV elimination,^6^ most HIV infected individuals must maintain a lifelong treatment regimen of ART and tolerate the toxic effects as well as considerable expense.^5^ To achieve a functional cure for HIV-1, it will be essential to enhance specific immune responses to control virus replication and eliminate virus-infected cells.^3^ It was appropriate to revisit anti-HIV CARs given the success of CARs for cancer immunotherapy, and many researches took advantage of novel CARs based on the new generation of bNAbs and showed the HIV-1-specific functional activity of those CARs.^17,18,19^

However, several immune checkpoint blockers showed upregulated expression in chronic HIV infection, such as PD-1 and Tim3, associated with the function decline and exhaustion of specific CTLs in HIV-infected individuals.^12,24,25,51^ Previous studies have tried to co-express the bNAb-based CARs and a PD-1 dominant negative receptor (DNR) that can saturate PD-1 ligands and thereby block signaling through the endogenous PD-1 receptor.^23^ In view of the potential efficacy of the blockade of PD-1 by using monoclonal antibodies for HIV cure, we first generated 3BNC117-E27 CAR-T cells to not only powerfully eliminate HIV-infected cells but also secret PD-1 blocking scFv E27 to overcome the immune inhibition caused by PD-1/PD-L1 pathway. Our work demonstrated that E27 could specifically bind to PD-1 on the surface of secreting 3BNC117-CAR T cells to achieve the blockade of PD-1 signaling, so as to enhance the anti-HIV potency of those CAR T cells with stronger proliferation capability, greater killing activity and enhanced cytokine secretion in the presence of target cells. We developed a platform for combining the application of anti-HIV CAR-T cells with the treatment of an anti-PD-1 blocker in a single step for adoptive T-cell therapies. Distinguished from DNR, which can only block the PD-1/PD-L1 pathway in CAR-T cells expressing DNR, E27 secreted from our 3BE CAR-T cells may affect much more other cells with high expression of PD-1 during HIV infection such as HIV-infected CD4+ T cells^52,53^ and play a broader role in the treatment of HIV. Furthermore, 3BE CAR-T cells can provide long-lasting expression and secretion of PD-1-specific antibodies, superior to one-dose injection of antibodies in view of their short half-life period. In addition, blockers targeting other immune checkpoint molecules like CTLA-4 and Tim3 can be taken into consideration in our platform for a potential strategy toward HIV functional cure.

Although the current CAR T-therapy showed promising results against HIV *in vitro* and in animal models,^54^ the autologous paradigm could be one of the various obstacles for the clinical application of CAR T-therapy. Personalized products might be challenging in patients with HIV-mediated immune deficiency due to lack of good quality T cells for engineering.^34^ Therefore, it is necessary to develop an approach to manufacturing “off-the-shelf” anti-HIV CAR-T cells from third-party donors. Since the key barrier to the adoptive transfer of third-party CAR T cells is the occurrence of GvHD, we took the first step to eliminate the endogenous TCR by CRISPR/Cas9 technology in anti-HIV CAR-T cells to avoid GvHD. Our approach could achieve high levels of the TCR-deficient anti-HIV CAR-T cells (the percentage of TCR-negative T cells could be over 97%), and it was proved that those T cells were failed to response to TCR stimulation *in vitro*, which suggested the functional inactivation of the TCR gene. A particularly significant aspect of our results was the demonstration of anti-HIV potency of those TCR-deficient 3BE CAR-T cells. Our studies showed that the functional activities of TCR-deficient anti-HIV CAR-T cells, such as specific killing activity and cytokine secretion, were equivalent to those of the conventional anti-HIV CAR-T cells. Those results suggested that the deficiency of TCR did not compromise their anti-virus potency relative to standard anti-HIV CAR-T cells, which could lay a solid foundation for the development of universal allogeneic anti-HIV CAR-T therapy. Further research is certainly needed to investigate whether those TCR-deficient 3BE CAR-T cells can powerfully kill HIV-infected cells *in vivo* without the occurrence of GvHD.

To sum up, we have provided a feasible approach to large-scale generation of “off-the-shelf” anti-HIV CAR-T cells with potent anti-virus activity and reduced alloreactivity by combing the lentiviral transduction of 3BE CAR with the electroporation of *TRAC*-4 RNP. This therapeutic strategy not only achieved a rational combination of CAR-T therapy and antibody therapy of PD-1 blockade for the treatment of HIV, but also provided universal anti-HIV CAR-T cells for adoptive transfer without a compromise of anti-virus potency. On account of the resource constraints of biological safety protection third-level laboratory at present, we have not thoroughly investigated the functions of 3BE/TCR-T cells *in vivo*. It will be necessary to extend our observations to a mouse model and further to HIV-infected individuals, to confirm the potential of our strategy to be a powerful therapeutic candidate for the functional cure of HIV.

## Materials and methods

### Anti-HIV CARs

The second-generation 3BNC1^17^ CAR was provided as the generous gift of Dr. Otto O. Yang.17 The PD-1 blocking scFv E27 gene was synthesized by GENEWIZ (GENEWIZ, Suzhou, China) according to the previous report.^36^ 3BNC117 CAR and E27 were cloned into a lentiviral vector (pTRPE-GFP-T2A-mRFP, as the generous gift of Professor James L. Riley) backbone downstream from an EF1a promoter.

### Cell lines

HEK293T cells and Jurkat cells were purchased from ATCC (Manassas, VA, United States). LHL2/3 cells and LEL6 cells were constructed in our lab and used somewhere else.^23^ LHL2/3 cells were HL2/3 cells (obtained through the NIH AIDS Reagent Program, Division of AIDS, NIAID, NIH: HL2/3 from Dr. Barbara K. Felber and Dr. George N. Pavlakis) encoded to highly express PD-L1 and firefly luciferase gene. LEL6 cells were Jurkat cells encoded to highly express Env, PD-L1 and firefly luciferase gene. HEK293T cells and LHL2/3 cells were maintained in DMEM (Gibco, Grand Island, NY, United States) medium supplied with 10% fetal bovine serum (FBS) (Gibco, Grand Island, NY, United States) and 1% penicillin-streptomycin (Gibco, Grand Island, NY, United States). Jurkat cells and LEL6 cells were maintained in RPMI1640 (Gibco, Grand Island, NY, United States) medium supplied with 10% fetal bovine serum (FBS) (Gibco, Grand Island, NY, United States) and 1% penicillin-streptomycin (Gibco, Grand Island, NY, United States).

### Isolation and culture of primary human T lymphocytes

Primary human CD3+ T cells were isolated from healthy donors, which from the Blood Center of Shanghai (Shanghai, China) and approved by the Ethics Committee of School of Life Sciences, Fudan University, by Ficoll-Paque gradient separation (GE Healthcare, Boston, MA, United States) and negative immunomagnetic bead selection according to the manufacturer’s protocol (Miltenyi Biotec, Germany). Primary lymphocytes were stimulated with CD3/CD28 T-cell activation Dynabeads (Gibco, Grand Island, NY, United States) for 48 hours. Those T cells were expanded in Vivo15 (Lonza, Basel, Switzerland) medium supplemented with 10% FBS and 5 ng/ml IL-2 (R&D, St. Paul, MN, United States) and 2 ng/ml IL-15 (R&D, St. Paul, MN, United States). Cell culture were maintained in an environment of 37°C and 5% CO2.

### Production of pseudoviruses and generation of anti-HIV CAR-T cells

To product pseudoviruses, HEK293T cells were seeded at 5 × 10^6^ cells per 10-cm dish. 14-16 hours later, pseudoviruses were generated by co-transfecting HEK293T cells with plasmids pTRPE encoding various CAR moieties (10 μg), psPAX2 (6.8 μg) and pMD.2G (3.4 μg) using PEI following the manufacturer’s instructions. Lentiviral supernatant was harvested 48 hours and 72 hours after transfection. Cell debris was removed by centrifugation at 5000 rpm for 20 min, followed by filtration through a 0.45 μM membrane (Millipore, Boston, MA, United States). Pseudoviruses were concentrated by centrifugation at 25000 rpm for 2 h at 4◦C, and then resuspended in the frozen stock solution and stored in −80◦C. For infection, 1 × 10^6^ stimulated CD3+ cells were transduced with concentrated pseudovirus supernatant at specific MOIs plus polybrene (Yeason) at 7 μg/ml for 12 hours, and then pseudoviruses were replaced by the fresh culture media as described above.

### Flow cytometry

The following monoclonal antibodies and reagents were used with the indicated specificity and the appropriate isotype controls. FITC-conjugated goat anti-human F(ab’)_2_ antibody (Jackson, Pennsylvania, United States), PE-conjugated mouse anti-human CD279 (PD-1) antibody (BioLegend, Santiago, CA, United States), FITC mouse anti-human TCRαβ (BD Biosciences, San Jose, CA, United States), FITC mouse anti-human CD25 (BD Biosciences, San Jose, CA, United States), PE mouse anti-human CD69 (BD Biosciences, San Jose, CA, United States), FITC mouse anti-human CD45RA (BD Biosciences, San Jose, CA, United States) and PE mouse anti-human CD62L (BD Biosciences, San Jose, CA, United States). All the cells were washed, resuspended in 100 μl PBS containing specific antibodies and incubated for 30 minutes at 4◦C. Data were acquired on a Beckman Coulter Gallios flow cytometer and analyzed by FlowJo version 10 (Tree Star, Ashland, OR).

### Western blotting

The supernatants were harvested and cells were lysed on ice for 30 minutes. The thermally denatured protein extracts were loaded on a 10% polyacrylamide gel, electroblotted onto a nitrocellulose membrane and blocked for one hour. The antibodies and dilutions used in these experiments were as follows: rabbit-anti-HA (1: 2500 dilution) (Abmart, Shanghai, China), rabbit anti-β-actin (1: 2500 dilution) (Proteintech, Wuhan, China), goat-anti-rabbit-secondary antibodies (1: 5000 dilution) (Beyotime, Shanghai, China). Bands were visualized using the ECL Western blotting system (Santa Cruz Biotechnology).

### Cytotoxicity Assay

Lactate dehydrogenase assay (LDH assay) was performed to measure the specific killing activity of anti-HIV CAR-T cells toward LHL2/3 cells and LEL6 cells at different ratios from 1:1 to 10:1 by using the CytoTox 96 non-radioactive cytotoxicity kit (Promega, Madison, WI, United States) following the manufacturer’s instructions. Briefly, anti-HIV CAR-T cells (effectors) and LHL2/3 cells or LEL6 cells (targets) were cultured together for 4-8 hours at various effector to target (E:T) ratios in a 96-well U-bottom plate at 37◦C. The supernatants were collected and incubated with 50 μL Cytotox 96 regent for 30 minutes. 50 μL Stop solution was added to each well and then absorbance values of wells were detected at 490 nm.

The expression level of luciferase was also detected after the coincubation at different E:T ratio to confirm the killing efficiency. Cells were harvested after the coincubation and resuspended in 50 μl PBS. Then, 50 μL Glo reagent (Promega) was added to each well and incubated for 5 minutes. Luminescence detection was performed by an inspired plate reader (Biotek, Montpelier, VT, United States).

### Proliferation and apoptosis assay

The proliferation of anti-HIV CAR-T cells was measured by using the Cell Counting Kit-8 (Dojindo Molecular technologies, Gaithersburg, MD, USA). Briefly, the CAR-T cells were incubated with LHL2/3 cells at 1:1 for 24 hours to 96 hours in a 96-well plate, then CAR-T cells were harvested into another 96-well plate and 10 μl of CCK-8 solution was added into each well of the 96-well plate and incubated for 1 h at 37°C. The optical density (OD) values were detected at 450 nm.

The apoptosis assays were performed using the PE Annexin V-apoptosis detection kit ◻ (BD Biosciences, San Jose, CA, United States) according to the manufacturer’s instruction. The cells were harvested and washed twice with cold PBS, and then resuspended in 1X Binding Buffer. Next, cells were stained by PE Annexin V and 7-AAD at room temperature (25°C) for 15 min in the dark. 400 μl of 1X Binding Buffer was added to each tube and the stained cells were analyzed by flow cytometry.

### Construction of CRISPRs and cell electroporation

CRISPR-Cas9 crRNA and tracrRNA were chemically synthesized by IDT (Coralville, Iowa, United States) and mixed at 1:1 followed by incubation at 95 °C for 5 min and then allowed to slowly cool to room temperature to provide annealed sgRNA. The following sgRNA targeting sequences were used in our study: *TRAC*-sgRNA-1: AGAGTCTCTCAGCTGGTACA; *TRAC*-sgRNA-2: TGTGCTAGACATGAGGTCTA; *TRAC*-sgRNA-3: ACAAAACTGTGCTAGACATG; *TRAC*-sgRNA-4: TCAGGGTTCTGGATATCTGT. S.p. HiFi Cas9 Nuclease V3 was purchased from IDT (Coralville, Iowa, United States). RNPs were produced by complexing sgRNA and spCas9 for 10 minutes at room temperature and should be electroporated immediately after complexing.

Electroporation was performed using Lonza 4D electroporation system (Basel, Switzerland) as the manufacturer’s instructions. For 20 μl Nucleocuvette Strips, 2 × 10^6^ cells resuspended in electroporation buffer were mixed with RNP complex (containing 80 pmol Cas9 and 80 pmol sgRNA) and transferred to a cuvette for electroporation. After electroporation, pre-warmed 100 ul media was added to the cuvette immediately, and cells were incubated at 37 °C for 15 min. Then, the electroporated cells were resuspended with pre-warmed media with FBS and cytokines as described above and transferred away from electroporation cuvettes.

### T7E1 assay and sequencing of PCR fragments

Cells were pelleted and lysed, and a sequence spanning the *TRAC* target site was PCR-amplified from cell lysates to be used to determine the level of genomic disruption of *TRAC* by a T7E1 Surveyor Nuclease assay (NEB, Ipswich, MA, United States). The PCR primers used for the amplification of the target locus were as follows: *TRAC* forward, 5’-TTCCCATGCCTGCCTTTACTC-3’; *TRAC* reverse, 5’-GCTGTTGTTGAAGGCGTTTG-3’. PCR products were ligated to pMD18-T cloning vector (Takara Bio, Beijing, China) and then transformed in E.coli. Single clone was picked and sequenced to calculate the indels and insertions.

### ELISA assays

Anti-HIV CAR-T cells or UTD cells were co-cultured with LEL6 cells (1 × 104 cells) at 10:1 for 24 hours in 96-U bottom well plates. Supernatants were collected and cytokines release by CAR-T cells in response to stimulation with target cells was analyzed using TNF-α and IFN-γ ELISA Kits (Dakewe, Shenzhen, China) according to manufacturer’s instructions.

### Statistics

Statistical analyses were performed using GraphPad Prism (GraphPad, San Diego, CA, United States). Experimental data are presented as mean ± SD and were analyzed by T-test or one-way ANOVA. A p value <0.05 was considered statistically significant.

## Acknowledgments

The authors thank Dr. Otto O. Yang, Professor James L. Riley, Dr. Barbara K. Felber, Dr. George N. Pavlakis, and Professor Edward A. Berger for providing materials.

## Author Contributions

Huanzhang Zhu was responsible for conceiving and designing of the whole study. Hanyu Pan performed most experiments and analyzed the data. Jing Wang and Huitong Liang participated in some of the experiments. Pengfei Wang kindly provided some suggestions for some experiences and revised the manuscript. The manuscript was prepared by Hanyu Pan and Huanzhang Zhu. All authors read and approved the submitted manuscript.

